# Determining epitope specificity of T cell receptors with TCRGP

**DOI:** 10.1101/542332

**Authors:** Emmi Jokinen, Jani Huuhtanen, Satu Mustjoki, Markus Heinonen, Harri Lähdesmäki

## Abstract

T cell receptors (TCRs) can recognize various pathogens and consequently start immune responses. TCRs can be sequenced from individuals and methods analyzing the specificity of the TCRs can help us better understand individuals’ immune status in different diseases. We have developed TCRGP, a novel Gaussian process method to predict if TCRs recognize certain epitopes. This method can utilize CDR sequences from TCR*α* and TCR*β* chains and learn which CDRs are important in recognizing different epitopes. We have experimented with with epitope-specific data against 29 epitopes and performed a comprehensive evaluation with existing prediction methods. On this data, TCRGP outperforms other state-of-the-art methods in epitope-specificity predictions. We also propose a novel analysis approach for combined single-cell RNA and TCR*αβ* (scRNA+TCR*αβ*) sequencing data by quantifying epitope-specific TCRs with TCRGP in phenotypes identified from scRNA-seq data. With this approach, we find HBV-epitope specific T cells and their transcriptomic states in hepatocellular carcinoma patients.

## Introduction

The adaptive immune system implements various complex mechanisms for surveillance against both pathogens and pathological cells arising in our body. To initiate an adequate adaptive immune response, a peptide, called epitope must first be bound by the major histocompatibility complex (MHC) class I or II molecule expressed on the surface of a nucleated cell or a professional antigen-presenting cell. The peptide-MHC (pMHC) complex is then presented to T cells which can recognize the complex via the T cell receptor (TCR) protein, consequently leading to T cell activation and proliferation by clonal expansion^1^. During clonal expansion, a fraction of T cells gain a long-living memory phenotype and therefore a clonal population of T cells with identical TCR rearrangements remain for years against the recognized antigen^2^, thus forming a potentially mappable immunological signature. Learning these signatures could have implications in broad range of clinical applications including infectious diseases, autoimmunity and tumor immunology.

T cells undergo non-homologous recombination during T cell development, which involves rearrangement of the germline TCR loci from a large collection of variable (V), diversity (D) and joining (J) gene segments as well as template-independent insertions and deletions at the V-D and D-J junctions^3,4^. TCRs are formed by a pair of *α* and *β*-chains (90-95% of T cells) or *γ* and *δ*-chains (5-10%) and V(D)J recombination happens in each locus independently. It is estimated that this rearrangement can result in the range of 10^18^ different TCR genes^5,6^ which provides enormous diversity for epitope-specific T cell repertoire. Furthermore, due to the low affinity of TCR-pMHC-interaction, TCR recognition is degenerate and a single TCR can interact with more than 1 million different epitopes (cross-reactivity), and a given epitope can elicit response from millions of TCRs^7,8^. Given these three levels of diversity, predicting TCR’s epitope specificity is notably challenging^9^.

The complementarity determining regions (CDRs) of a TCR determine whether the TCR recognizes and binds to an antigen or not^10^. Of these regions, CDR3 is the most variable and primarily interacts with the peptide, while CDR1 and CDR2 primarily interact with the peptide binding groove of the MHC protein presenting the peptide, but can also be in contact with the peptide^11,12^. Dash et *al*.^13^ noted that also a loop between CDR2 and CDR3 (IMGT^®14^ positions 81-86), which they called CDR2.5, has sometimes been observed to make contact with pMHC in solved structures. Figure 1 shows these CDRs in interaction with a peptide-MHC-complex (pMHC).

**Figure 1:**
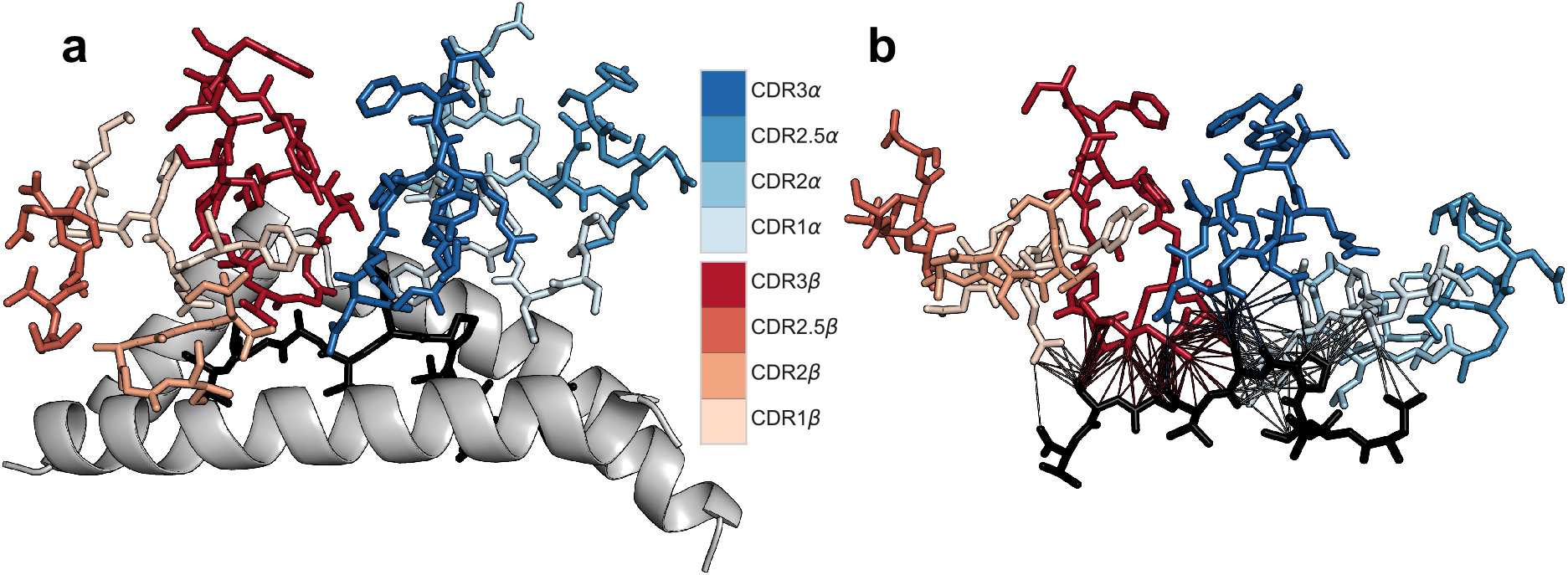
Structure of a TCR-pMHC-complex. **a)** CDRs of a TCR binding to CMV-epitope pp65_495-503_ (shown in black), presented by MHC-protein, whose binding groove is shown as a white cartoon. **b)** Distances below 5 Å between the atoms of the epitope and the different CDRs are shown. The original structure was determined by Gras et *al*.^15^.

It is well known that for example the CDR3*β* of a TCR is important in recognizing peptides presented to the T cell, but it is still unclear which specific physicochemical or structural features of the CDR3*β* or of other parts of the TCR determine the antigen recognition specificity of that T cell. Although high-throughput DNA sequencing has enabled large-scale characterization of TCR sequences^6^, it still remains exhaustive to profile epitope-specific TCRs as they require sample-consuming experiments with distinct pMHC-multimers for each epitope of interest. Therefore, there is a great need for models that examine which epitopes a TCR can recognize or to which TCRs an epitope can bind to. Curated databases of experimentally verified TCR-peptide interactions have recently been launched, such as the VDJdb, IEDB, and McPAS^16,17,18^. Such data sources are enabling more comprehensive, data-driven analysis of TCR-peptide interactions, and make it possible to use techniques from statistical machine learning for the aforementioned tasks. Yet only a few computational TCR specificity models have been proposed in the literature^12,13,19^, some of which rely on heuristics, may be suboptimal for small datasets and have not been benchmarked against each other.

We propose a method called TCRGP which builds on non-parametric modelling using Gaussian process (GP) classification. Probabilistic formulation of GPs allows robust model inference also from small data sets, as is currently the case for TCR-peptide interaction information in curated databases. As the space of all TCRs that can recognize a certain epitope is potentially very large, it is important to avoid overfitting to the limited sample of TCRs that is available. Indeed, TCRGP clearly outperforms the current state-of-the art methods for predicting the epitope specificity of TCRs. At the same time, TCRGP also scales to extremely large data sets which we expect for the future epitope specific TCR-seq data sets. We also analyze the effects of utilizing different sections of the TCR amino acid sequence, and examine how the number of available TCRs for training affects the predictions. Finally, we demonstrate the usefulness of TCRGP in analyzing single-cell RNA+TCR*αβ*-sequencing data from hepatocellular carsinoma patients.

## Results

### Gaussian process classifier for TCRs

Gaussian processes (GP) are a flexible class of models that have become popular in machine learning and statistics with various applications in molecular biology, bioinformatics and other fields^20,21,22,23,24^. We have developed TCRGP, a Gaussian process based probabilistic classifier to predict TCRs’ epitope specificity. GPs are nonparametric and differ from standard parametric models in that they define priors for entire nonlinear functions, instead of their parameters. GPs implement a Bayesian nonparametric kernel method for learning from data. Properties of GPs are defined by the kernel function, which is a function of objects that we want to classify. Our objects are amino acid sequences (strings) that have varying lengths. While kernel functions can be defined for strings, we use a feature representation that first aligns the amino acid sequences into a fixed length presentation using IMGT^®^ definitions. BLOSUM based substitution matrices are then used to measure similarities between aligned amino acids via squared exponential kernel function, whose hyperparameters control the complexity and smoothness of the classifier.

TCRGP utilizes GPs’ probabilistic nature and infers the classifier using variational inference. Probabilistic inference makes the method more robust in small data regime, where the current experimental data sets are, while sparse variational inference scales the method to extremely large (future) data sets. Since it is currently poorly understood that how different CDR regions and *α*/*β* chains (features) contribute to the epitope specificity (see Fig. 1), we extend TCRGP to use all these features by using multiple kernel learning and use experimental data to automatically calibrate the strength of each feature’s contribution to the final classifier. See Methods Section for a detailed description of TCRGP method.

We use two data sets to demonstrate TCRGP’s accuracy in predicting TCR epitope specificity: a recently published data set of tetramer sorted TCR sequences for 10 epitopes, introduced by Dash et *al*.^13^, and a new dataset of medium and high quality epitope-specific TCR sequences extracted from VDJdb database^16^. The Dash data provides the largest set of epitope-specific paired TCR*αβ*-data that we are aware of, and VDJdb provides a comprehensive selection of available epitope-specific TCRb-data currently available. We also considered using using TCRs from IEDB^17^ and McPAS^18^, but they had significant overlap with VDJdb and their collections of TCRbs were not as extensive. Both of the selected data sets are combined with a set of background TCRs, also presented by Dash et *al*.^13^ that are not expected to recognize the epitopes in the two data sets. See Methods Section for details for the data sets. Our work is accompanied by an efficient software implementation that contains pre-existing models for predicting TCRs’ specificity to epitopes involved in data sets used in this study as well as tools for building new epitope specificity models from new datasets.

### Evaluating the significance of utilizing different CDRs

To evaluate the benefit of using different CDRs, we used the data set of Dash et *al*.^13^ which includes 4635 pMHC-tetramer sorted single-cell sequenced TCR*αβ* clonotypes from 10 epitope-specific repertoires (from hereon referred as the Dash data). We trained our TCRGP model using either only CDR3 or also with CDR1, CDR2, and CDR2.5 from TCR*α*, TCR*β*, or both. We applied leave-one-subject-out cross-validation as described in Methods Section. Figure 2 a presents the cross-validation results for a single BMLF1_280-288_-epitope from EBV and demonstrates how the classification results vary between different subjects likely due to the high variety of the TCRs.

**Figure 2:**
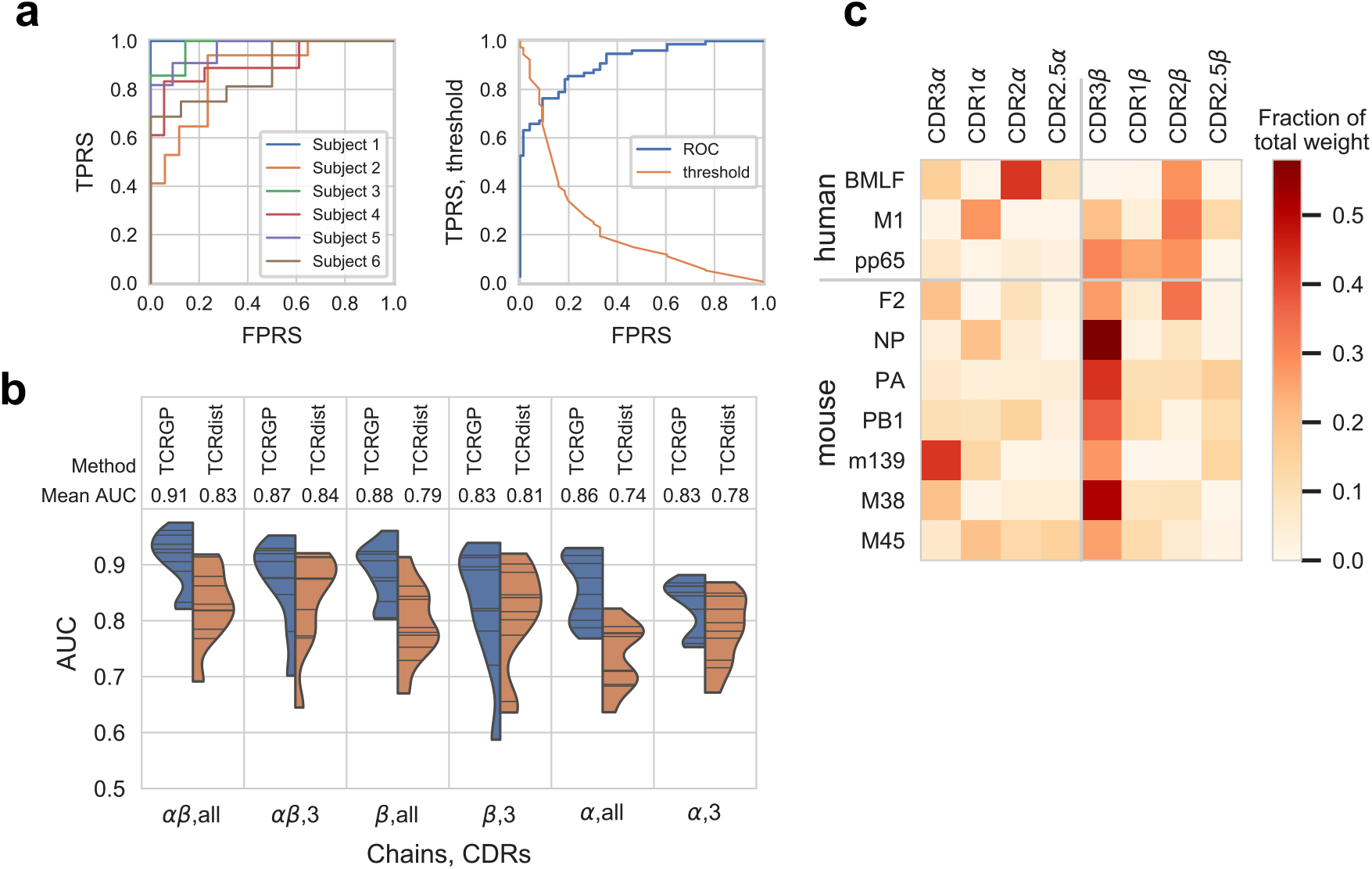
Epitope-specificity prediction with Dash data. **a)** The left panel shows the cross-validated ROC curves for each subject in the Dash data for BMLF1_280-288_, when TCRGP has been trained using all CDRs from TCR*α* and TCR*β*. The mean AUROC is 0.905. The right panel shows the ROC curves when the predictions have been combined and also the corresponding threshold values. From this figure we can determine which threshold values correspond to different true positive rates (TPRS) and false positive rates (FPRS). **b)** The blue parts of the violin plots illustrate the AUROC-scores of predictions made by TCRGP for all the epitopes. The orange sides illustrate the AUROC-scores obtained with TCRdist. A horizontal line within a violin plot presents the mean AUROC-score obtained for one epitope. The used chains (*α* and/or *β*) and CDRs (three or all) are indicated below each panel. **c)** Fractions of total weight given to kernels corresponding to different CDRs, when TCRGP has been trained to predict which TCRs are specific to the epitopes in the Dash data using all CDRs from both TCR chains.

AUROC-scores of the predictions for different combinations of CDRs and *α*/*β* chains are summarized in Fig. 2 b. For comparison, we also trained TCRdist in the same manner. Figure 2 b shows that both methods, TCRGP and TCRdist, perform on average better when using TCR*β* than when using TCR*α*, although using both *α* and *β* chains generally provides the best results. There are few exceptions, as shown in Supplementary Fig. S1. For example, with peptides pp65 both models perform better when using CDR3*α* instead of CDR3*β*. Overall TCRGP is better than TCRdist in utilizing information from CDRs other than CDR3. TCRGP achieves higher AUROC-scores on average when trained using all CDRs instead of only CDR3, whereas with TCRdist the AUROC-scores seem to be similar or better when only CDR3 is utilized. Notably TCRGP outperforms TCRdist in prediction accuracy for 57 of the 60 comparisons (Supplementary Fig. S1, Fig. 2 a).

Figures 2 b and S1 also show that the AUROC-scores can have notable differences between different epitopes even when the same combinations of CDRs and *α*/*β* chains have been utilized. Some of these differences may be explained by the differences in the number of available training samples, for example for pp65 there were only 76 TCRs from 6 subjects in the Dash data, which may have contributed to a lower prediction accuracy. To address this, we evaluated the models also using leave-one-out cross-validation with only unique, private TCRs to see how the models perform when predictions are done only on new TCRs. We consider a TCR to be unique when it consists of a unique combination of CDR3 amino acid sequence and V-genes from both chains. With both TCRGP and TCRdist, the average AUROC-scores improve slightly (Supplementary Fig. 2 and Fig. 3), demonstrating that the models can predict the specificity of completely new sequences and that the larger number of TCRs used for training (due to the larger folds in leave-one-out cross validation) improve the model performances.

**Figure 3:**
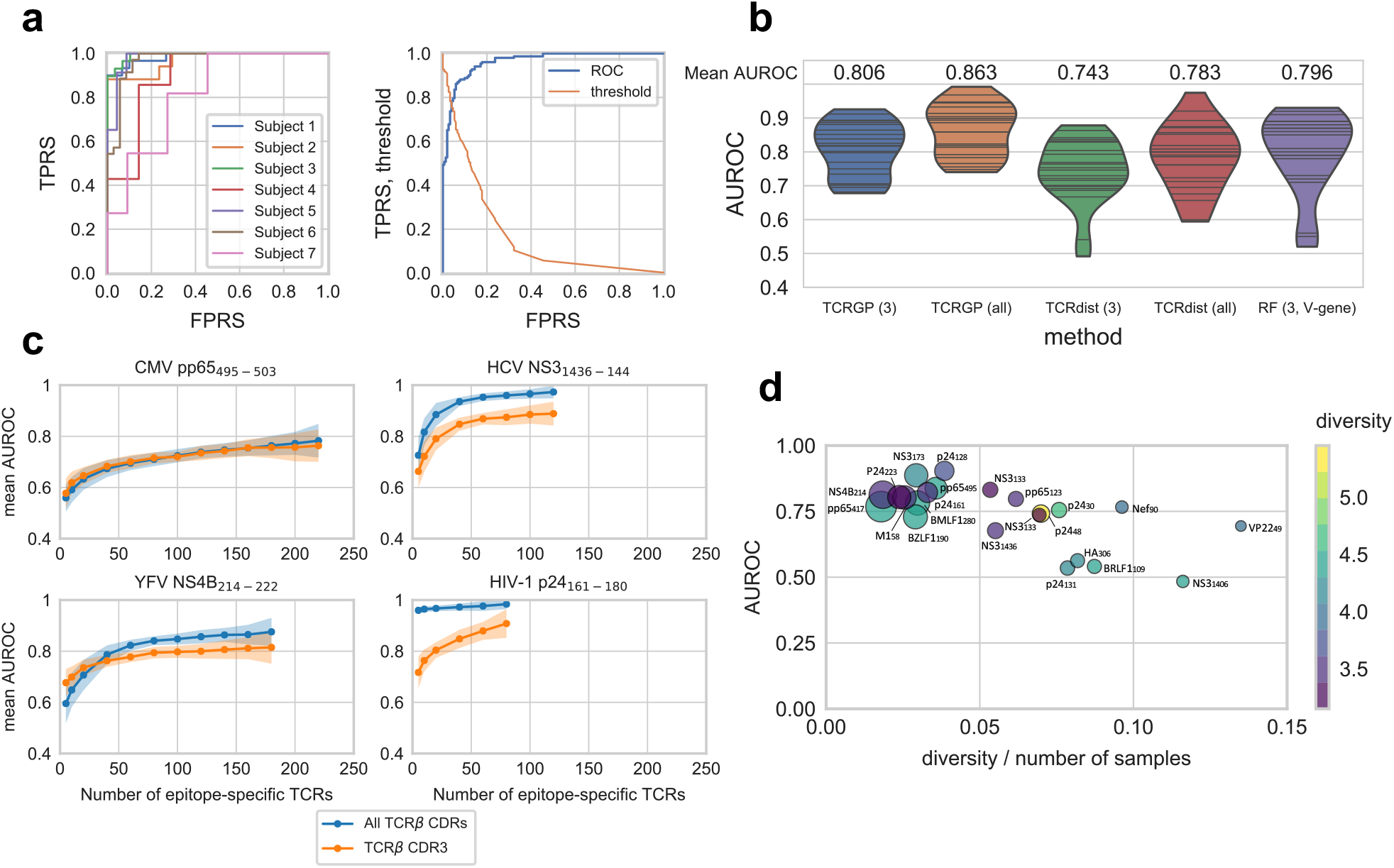
Epitope specificity prediction with VDJdb data. **a)** Left panel shows the cross-validated ROC curves for each subject in the VDJdb data for HCV NS3_1436-1444_-epitope, when TCRGP has been trained using TCR*α* and TCRβ with all CDRs. The mean AUROC is 0.944. Right panel shows the ROC curves when all predictions have been combined and also the threshold values for classification are shown. From this figure we can determine which threshold values correspond to different true positive rates (TPRS) and false positive rates (FPRS). **b)** One violin plot presents the mean AUROC-scores obtained with one method for all epitopes in our VDJdb data. Below each violin plot there is the name of the method used and in the brackets which CDRs have been used (3 for CDR3, all for CDR1, CDR2, CDR2.5, and CDR3). A horizontal line within a violin plot presents the mean AUROC-score obtained for one epitope. RF refers to the Random Forest TCR-classifier of De Neuter et *al*.^19^. **c)** For each epitope from the VDJdb dataset, TCRGP models were trained using different numbers of unique epitope-specific TCRβs, always complemented with the same number of control TCRβs. For each point of the learning curve the model was trained with 100 random samples of the TCRβs, using either CDR1, CDR2, CDR2.5, and CDR3 (blue curves), or only CDR3 (orange curves). The darker lines show the mean of the predictions and the shaded areas ± the standard deviation for the 100 folds. The points indicate the tested sample sizes. Here learning curves for four peptides are shown, **d)** Leave-one-out cross-validated AUROC-scores correlate with the diversity and number of samples (Pearson correlation −0.66). The sizes of the circles indicate the number of unique TCRs used for training (see Table 3).

To better understand the significance of the different CDRs for TCR-pMHC recognition, we also examined more closely how TCRGP weighted the kernels created for the different CDRs, when all CDRs from both chains were utilized. Figure 2 c illustrates which CDRs were found important for the different epitopes. As one might expect, with most of the epitopes most weight was given to the CDR3*β*, but utilizing several epitopes was found beneficial with all epitopes. This is in agreement with an alignment of 52 TCR sequences from TCR-pMHC PDB structure complexes, which demonstrates that all CDRs can be within 5Å of peptide^12^. For example with CMV-epitope pp65_495–503_, experimental characterization of the structure observed contacts between the peptide and CDR3*β*, CDR1*β*, CDR3*α* and CDR1*α* (Fig. 1 b), and CDR2*β* is also in proximity of the peptide (within 5.8Å). Another TCR-pMHC-complex structure (PDBid 5D2L) for the same pp65_495–503_-epitope suggests that CDR2*β* was also within 5 Å of the peptide. Indeed, the optimized weights for the pp65 epitope (Fig. 2 c) show some correspondence to the observed contacts (Fig. 1 b). However, with some epitopes CDR3*β* was not considered very important, as for example with mCMV-epitope m139_419-426_ CDR3*α* is more important for the prediction, while with EBV-epitope BMLF1_280-288_ most of the weight was given to CDR2*α* and CDR2*β*.

### Comparisons to other methods

We also experimented with a data set we obtained from VDJdb, which gathers published epitope-specific TCR-sequencing results and is currently the largest collection of such data (from hereon referred as the VDJdb data). We again used leave-one-subject-out cross-validation as described in Methods Section. We trained TCRGP and TCRdist using only CDR3*β* and then also with the other CDR*β*s. Figure 3 a shows the ROC curves when TCRGP was trained with all CDR*β*s to predict which TCRs are specific to the HCV-epitope NS3_1436–1444_.

We also trained a TCR-classifier as proposed by De Neuter et *al*.^19 19^ using the same data. Unfortunately, the background TCR data set from^13^ did not contain information of the J-gene, which is requested by this TCR-classifier. However, according De Neuter et *al*.^19^ themselves, not much weight was given to the J-gene at least in their experiments. This TCR-classifier does not utilize other CDR*β*s in addition to CDR3*β*, but the V*β*-gene from which the CDR*β*s can be derived from. Thus all these three methods get the same sequence information, when TCRGP and TCRdist use all CDRs, although in a slightly different form.

Figure 3 b shows the distributions of mean AUROC scores for each model trained for the 22 different epitopes. With this data set from VDJdb, we can see that TCRGP and TCRdist both perform better, when all CDR*β*s have been utilized. Remarkably, TCRGP outperforms the other methods when using all CDRs, but also when only the CDR3*β* is utilized. AUROC scores for the different epitopes are presented in Supplementary Fig. S4.

In the VDJdb data, there were also TCRs that appeared in samples collected from multiple subjects (see Table 3). We therefore trained the models also using leave-one-out cross-validation with only unique TCRs. In this case we considered a TCR to be unique if it had a unique combination of CDR3*β* amino acid sequence and V*β*-gene, as we only utilized the TCR*β*. As with the Dash data above, our results (Supplementary Fig. S5 and Fig. S6) show that the models can predict the specificity of completely new sequences, thus demonstrating their use for epitope specificity prediction for previously unseen TCRs.

### Significance of the number of training samples

To assess how the number of epitope-specific TCRs affects the performance of TCRGP classifier, we trained our model using different numbers of epitope-specific TCRs from the VDJdb data. We selected all unique TCRs for each epitope and took 100 random samples from them for each training set size, using always an equal number of randomly chosen control TCRs. Learning curves for four epitopes are shown in Fig. 3 c and learning curves for all 22 epitopes in Supplementary Fig. S7. In general, the predictive performance of the TCRGP classifiers improve when more training samples are available. However, the exact number of TCRs required to achieve a certain level of accuracy varies greatly between the different epitopes. This likely reflects the fact that different epitopes can be more selective in choosing their TCR interactions. In other words, TCRs that recognize one epitope can be more diverse than the TCRs that recognize another epitope^13^, and if the TCRs are very heterogeneous, it requires more sampling to get a representative sample of these TCRs for the model training. Indeed, we observed a negative correlation between TCRs’ diversity and prediction accuracy (Fig. 3 d).

These learning curves also further demonstrate the benefit of using multiple CDR sequences: With most of the epitopes using all CDRs produces better or comparable AUROC-scores with all sample sizes, although there are a few epitopes with which the AUROC-scores are higher when utilizing only the CDR3*β* if the sample sizes are very small (≤ 40). These results also suggest that with many epitopes it may be more beneficial to sequence a moderate amount of TCRs in such precision that in addition to the CDR3 also the V-gene and allele (and thus the CDR1, CDR2, and CDR2.5) can be determined, than to sequence large amounts of only CDR3s. These findings are inline with TCRGP-predicted weights for each CDR3 for individual epitope, as we can see in the case of CMV-epitope pp65_495-503_, EBV-epitope BMLF1_280–288_ and IAV-epitope M1_58–66_. With pp65_495-503_ most weight was given to CDR3/3 and thus information from other CDR3s are not as beneficial; with BMLF1_280–288_ almost no weight was given to CDR3/3 and in the learning curves there is a clear improvement when all CDR/3s are used; with M1_58–66_ some weight was given to CDR3/3, but most weight fell to CDR2β and correspondingly there is a small improvement in the learning curves, when all CDR/3s are utilized. Overall, the learning curves show that TCRGP can learn an accurate predictor even from a small data set, thus making it applicable to the currently existing TCR-peptide interaction data sets. On the other hand, our results also show that TCRGP’s prediction accuracy increases along with increasing number of training examples, enabling analysis of larger TCR-peptide interaction data sets in the future.

### Leveraging TCRGP in single-cell RNA+TCR*αβ*-sequencing data analysis

We next demonstrate how TCRGP can be utilized to implement a novel analysis of combined single-cell RNA and TCR*αβ* (scRNA+TCR*αβ*) sequencing data. Hepatocellular carcinoma (HCC) is one of the leading causes for cancer-related deaths worldwide^25^. Globally the predominant cause of HCC is considered to be Hepatitis B virus (HBV) as half of the HCC patients are estimated to be chronic HBV carriers^26^. During the course of natural infection, HBV integrates itself into the genome of the hepatocytes and thus a proportion of the HCC cells expresses HBV antigens^27^. Therefore, the malignant cells could be targeted by HBV-specific T-cell clonotypes and the high-dimensional characterization of these clonotypes could be crucial in understanding the viral control of HBV-infection and its association to HCC. To address this previously unanswered question we used TCRGP to analyze a published single-cell RNA and TCR*αβ* dataset by Zheng et *al*.^28^ of T cells from HBsAg-positive HCC-patients from blood, non-malignant liver tissue and tumour tissue (from hereon referred as the Zheng data).

Recently, Cheng et *al*.^29^ mapped HBV-reactive T cell populations by exhaustively screening the whole HBV genome with an HLA-class I restricted multiplexed pMHC-tetramer strategy and characterized T cells against four interesting HBV-epitopes from two antigens with TCR*β*-sequencing. We used this TCR*β* data to train TCRGP classifiers (see Methods section for details) to enable prediction for the unselected TCR repertoire in the Zheng data against the four epitopes (HBV_core169_, HBV_core195_, HBV_pol282_, HBV_pol387_, where core refers to core protein and pol to polymerase protein) (Fig. 4 a).

**Figure 4:**
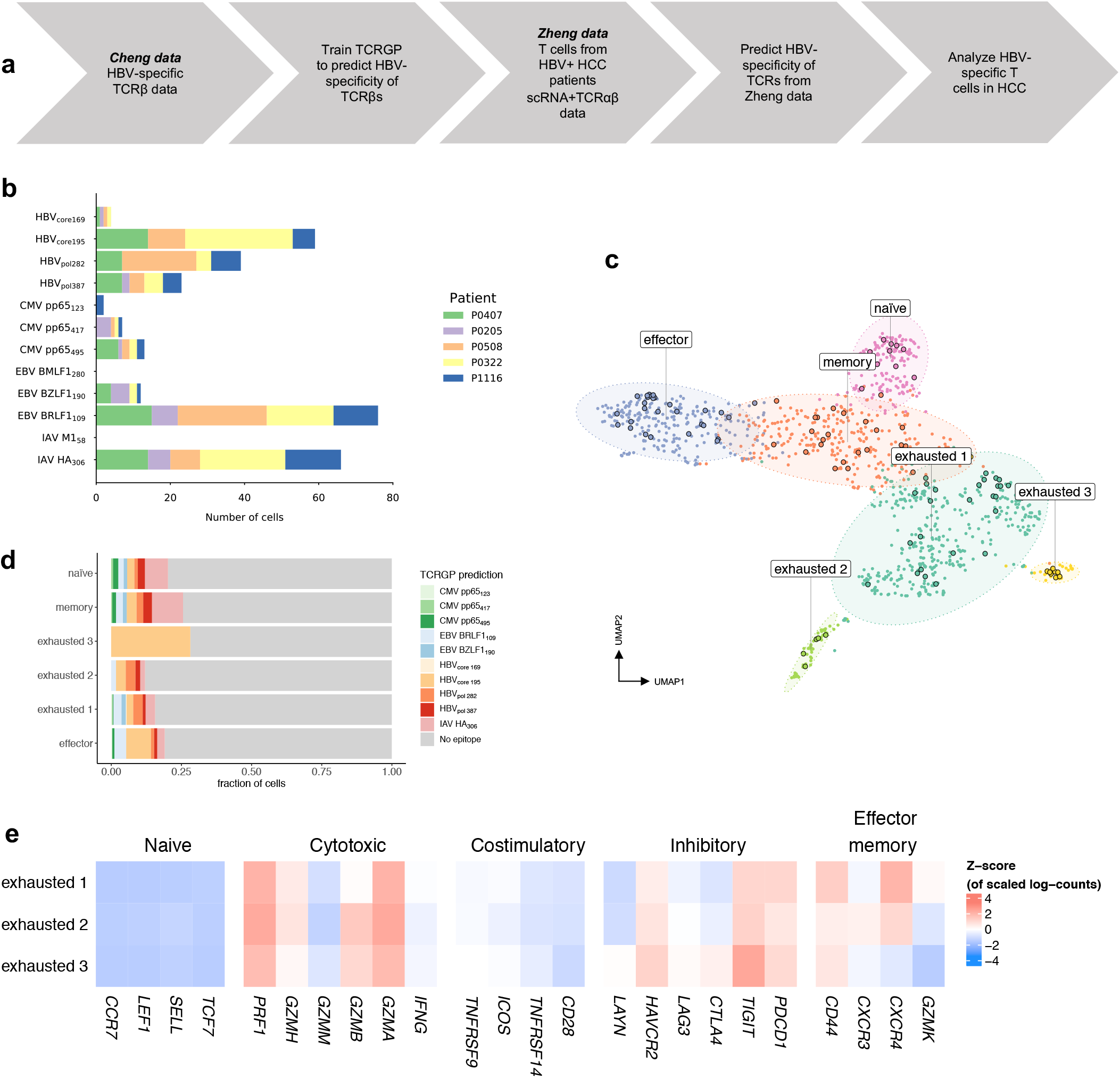
Analysis of HBV-specific T cells in HCC patients. **a)** Schematics for the analysis of single-cell RNA and TCR*αβ* sequencing data using TCRGP and multimer-sorted data, **b)** Numbers of cells predicted to recognize different epitopes by TCRGP with probability of at least 85%. HBV-reactivity was assessed by four different TCRGP classifiers trained against four different HBV-epitopes (HBV_core169_, HBV_core195_, HBV_pol282_, HBV_pol387_). Other predictions were made using the models trained with the VDJdb data, **c)** Dimensionality reduced representation (UMAP) of the 1189 CD8+ T cells from HBV+ HCC-patients from peripheral blood, normal adjacent tissue and tumour tissue. Encircled dots represent the T cells predicted to be HBV-reaetive by TCRGP. **d)** The frequencies of T cells predicted to recognize different HBV-epitopes in each cluster, **e)** The frequencies of T cells predicted to recognize different viral epitopes in each cluster, **f)** Z-score normalized mean expressions of known canonical markers to assess CD8+ cell phenotypes (naïve, cytotoxic, costimulatory inhibitory, and effector memory markers) in the three different exhausted cell clusters. Exhausted 3 was predicted to be enriched for HBV-targeting T cells (p=2.913e-06, p.adj=0.00103).

Of the 789 CD8+ cells from Zheng data analyzed with TCRGP, 108 cells (13.688%), were predicted to be reactive against HBV with at least a probability of 85%, most of which against HBVcore195-epitope (59 cells) (Fig. 4 a, b and c). On the contrary, 176 cells were predicted to be reactive against common viruses (CMV=22, EBV=88 and Influenza A=66 cells) (Fig. 4 b), showing that HBV was the most common target for antigen-specific T cells in HCC patients.

After unsupervised clustering of the CD8+ cells’ scRNA-seq data, we received 6 different phenotypes that were similar to the phenotypes described by Zheng et *al*.^28^, but had the exhausted cells divided into 3 different clusters instead of one (naïve, effector, memory, exhausted 1, exhausted 2 and exhausted 3) (Fig. 4 c). Interestingly, cells in exhausted 3 cluster showed the highest enrichment of the clonotypes targeting HBV_core195_-epitope (Fisher’s exact test p=2.913e-06, Benjamini-Hochberg corrected for multiple testing p.adj=0.00103), but not to any other epitope-specific clonotypes (Fig. 4 d, e). By calculating exhaustion score for each T cell, we found that exhausted 3 cluster was the most exhausted (against exhausted 2 p=0.0032, against exhausted 1 p=0.0021) and the least cytotoxic cluster (p=0.019 and p=1.3e-05). Further, gene-level analysis showed high expression of *TIGIT* and *HAVCR2* (encoding TIM-3), which have been associated with late-stage exhaustion after long antigen exposure. Upregulated pathways for exhaustion cluster 3 were IL2-STAT5 signaling pathway (exh3 vs exh1 q=0.022 and exh3 vs exh2 q=0.000) and myogenesis pathway (q =0.016 and q=0.001).

In summary, TCRGP was able to identify a T cell cluster that was enriched with HBV-targeting clonotypes, which was the most exhausted and least functional. These differences in transcriptomes of the exhausted clonotypes could explain the role of viral control in the development of HCC in HBV-carriers, which is further complicated by differential expression of HBV antigens that can elicit either more effector or exhaustion-prone immune response.

## Discussion

In this paper we have demonstrated that we can accurately predict if a previously unseen TCR can recognize an epitope, for which we have had sufficient amount of experimentally produced epitope-specific TCR-sequencing data for training. The performances of the models for different epitopes depend greatly on the size, quality, and heterogeneity of the repertoire of the epitope-specific T cells available for training, and not all the epitopes elicit oligoclonal responses that can be interpreted with machine learning models. We have also shown that the other CDRs in addition to CDR3*β* can provide useful information for the classification task and that it can depend on the epitope in question which of the CDRs are important.

As the amount of epitope-specific T cell sequences has expanded recently and the computational methods available are fairly new, no comparative effort has thus far emerged to gain understanding of the already available toolbox. In this work we provide the most thorough analysis of the current epitope-specificity prediction algorithms on the biggest data sets publicly available and thus provide important information to the community. The currently available sequence data of epitope-specific TCRs has allowed us to come this far, but it will be interesting to see what can be achieved when more data becomes available with modern high-throughput techniques presented recently^30,31^. Because of the limited data, we have not yet been able to consider the similarities of the epitopes and the significance of the different HLA-types of the MHC proteins presenting the epitopes. Having a larger variety of epitopes and TCRs that recognize them, would allow us to also better model the cross-reactivity of the TCRs. Eventually, when larger dataset become available, we may be able to model the similarity of the epitopes and consider the HLA-types in addition to the similarity of the TCRs and even predict if a TCR can recognize a previously unseen epitope. Furthermore, the proposed Gaussian process formalism has been shown to scale to very large datasets up to billion data points^32^ by optimizing a small number of landmark sequences.

The previous supervised algorithms developed are presented in the case of epitope-specific data, but we believe that to answer clinically relevant questions we need to address the unselected repertoire data which is far more numerous in size and more easily produced. Therefore we presented a novel workflow for analysis of scRNA+TCRab data in a clinically relevant question, showing the power of determining the epitope-specifity *in silico* to reveal underlying transcriptomic heterogeneity of the epitope-specific T cells, which to our knowledge has not been tackled before with single-cell RNA-sequencing in tumor infiltrating lymphocytes in any tumor. As the number of scRNA+TCRab and conventional TCRb sequencing data in clinical settings is increasing^33,34,35,36,37,38,39,40^, we expect that models like ours can be applied to a variety of research questions where exhaustive *ex vivo* pMHC-multimer assays are not feasible. In conclusion, we propose that TCRGP could be useful in the diagnosis and follow-up of infectious diseases, in autoimmune disorders and cancer immunotherapy.

## Methods

### Data

Our experiments focus on TCRs formed by a pair of *α* and *β* chains, as those are the most common type of TCRs^41^. The CDR3 sequence is formed by V(D)J recombination, but CDR1, CDR2, and CDR2.5 sequences are determined completely by the V-gene and allele^3^. Dash et *al*.^13^ provide a table of all V-gene and allele combinations and the corresponding CDR1, CDR2, and CDR2.5 amino acid sequences aligned according to IMGT^®^ definitions^14^. Our method can utilize the aligned amino acid sequences of all these CDRs from either one or both of the *α*- and *β*-chains of the TCR. Table 1 shows a few examples of TCR sequences and their alignment.

**Table 1:**
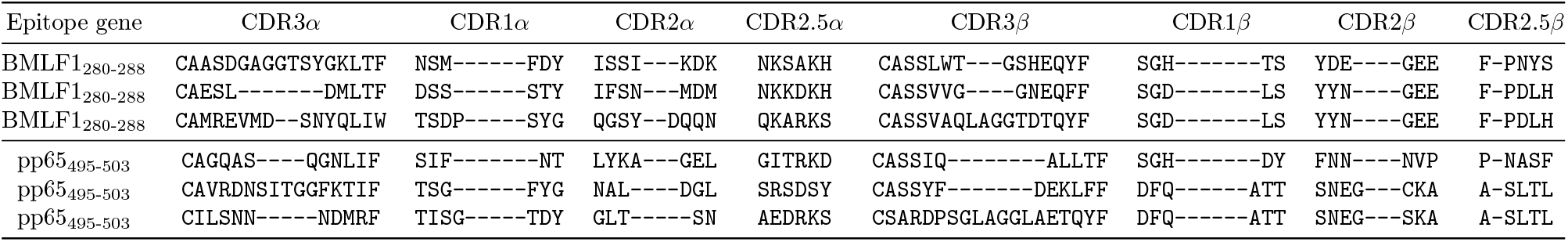
An example of aligned TCR sequences (from the Dash data) for two peptides. Each CDR type has been aligned separately according to IMGT definitions. CDR1, CDR2 and CDR2.5 sequences for both the *α*- and *β*-chains are defined by germline V*α*- and V*β*-genes. The alignments for all possible CDR1, CDR2 and CDR2.5 sequences have been determined by Lefranc^14^ and we use these alignments with all epitopes. CDR3s are aligned by adding a gap at the top of the loop^14^. The length of the alignment can then be determined by the length of the longest CDR3 available and can vary between different models.

In our experiments, we use a data set collected by Dash et *al*.^13^, which contains epitope-specific paired TCR*α* and TCR*β* chains for three epitopes from humans and for seven epitopes from mice, see Table 2 for details.

**Table 2:**
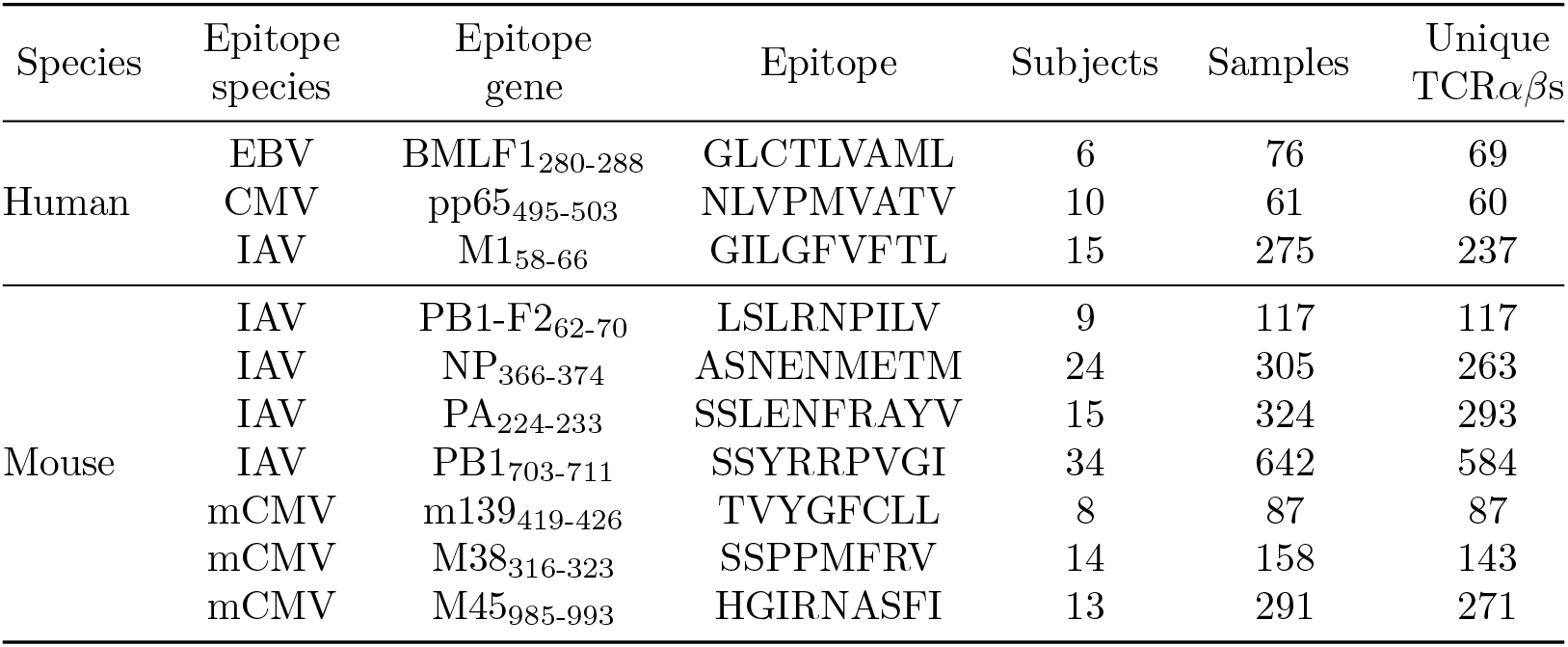
The Dash data contains epitope-spcefic TCRs for Epstein-Barr virus (EBV), human Cytomegalovirus (CMV), Influenza A virus (IAV) and mouse Cytomegalovirus (mCMV).

We also gather a new data set from VDJdb (https://vdjdb.cdr3.net), which is a database that contains TCR sequences with known antigen specificity^16^. Every entry in VDJdb has been given a confidence score between 0 and 3 (0: critical information missing, 1: medium confidence, 2: high confidence, 3: very high confidence). We constructed our data set so that we selected all epitopes that have at least 50 TCR*β* sequences with a confidence score at least 1 and found 22 such epitopes, see Table 3 for details. VDJdb also contains TCR*α* sequences, but since these are not in general paired with corresponding TCR*β* sequences, we chose to only experiment with the TCR*β* sequences.

**Table 3:** Data set gathered from VDJdb contains epitope-specific TCRs for Cytomegalovirus (CMV), Epstein-Barr virus (EBV), Influenza A virus (IAV), Hepatitis C virus (HCV), Herpes Simplex virus type 2 (HSV-2), Yellow Fever virus (YFV), Dengue virus type 1 (DENV1), Dengue virus type 3 (DENV3-4), and Human immunodeficiency virus type 1 (HIV-1).

| Epitope species | Epitope gene | Epitope | Subjects | Samples | Unique TCRβs |
| --- | --- | --- | --- | --- | --- |
| CMV | pp65<sub>123-131</sub> | IPSINVHHY | 17 | 65 | 58 |
| CMV | pp65<sub>417-426</sub> | TPRVTGGGAM | 29 | 184 | 122 |
| CMV | pp65<sub>495-503</sub> | NLVPMVATV | 103 | 413 | 242 |
| EBV | BMLF1<sub>280-288</sub> | GLCTLVAML | 54 | 299 | 152 |
| EBV | BZLF1<sub>190-197</sub> | RAKFKQLL | 17 | 225 | 149 |
| EBV | BRLF1<sub>109-117</sub> | YVLDHLIVV | 6 | 66 | 51 |
| IAV | M1<sub>58-66</sub> | GILGFVFTL | 50 | 239 | 138 |
| IAV | HA<sub>306-318</sub> | PKYVKQNTLKLAT | 11 | 56 | 50 |
| HCV | NS3<sub>1073-1081</sub> | CINGVCWTV | 7 | 76 | 39 |
| HCV | NS3<sub>1406-1415</sub> | KLVALGINAV | 4 | 65 | 65 |
| HCV | NS3<sub>1436-144</sub> | ATDALMTGY | 7 | 152 | 139 |
| HSV-2 | VP22<sub>49-57</sub> | RPRGEVRFL | 5 | 68 | 29 |
| YFV | NS4B<sub>214-222</sub> | LLWNGPMAV | 5 | 223 | 198 |
| DENV1 | NS3<sub>133-142</sub> | GTSGSPIVNR | 11 | 65 | 59 |
| DENV3-4 | NS3<sub>133-142</sub> | GTSGSPIINR | 8 | 51 | 46 |
| HIV-1 | p24<sub>30-40</sub> | KAFSPEVIPMF | 44 | 134 | 104 |
| HIV-1 | p24<sub>48-56</sub> | TPQDLNTML | 21 | 52 | 40 |
| HIV-1 | p24<sub>128-135</sub> | EIYKRWII | 12 | 81 | 60 |
| HIV-1 | p24<sub>131-140</sub> | KRWIILGLNK | 27 | 212 | 141 |
| HIV-1 | p24<sub>161-180</sub> | FRDYVDRFYKTLRAEQASQE | 17 | 141 | 95 |
| HIV-1 | p24<sub>223-231</sub> | GPGHKARVL | 1 | 62 | 53 |
| HIV-1 | Nef<sub>90-97</sub> | FLKEKGGL | 21 | 104 | 78 |

For the training and testing of the models, we also required some background TCRs that we do not expect to recognize the epitopes in our data sets. For this purpose we used a set of background TCRs constructed in Dash et *al*.^13^.

The data sets we have used can be found from github.com/emmijokinen/TCRGP.

### Leave-one-subject-out cross-validations

For the evaluation of the methods developed by us and others, we needed to divide our data sets for training and testing. Both of the data sets we use determine the subjects from whom each TCR in the data has been obtained from. We therefore chose to use leave-one-subject-out cross-validation, where we leave out all TCRs from one subject, train the model with all the other TCRs, test it with the TCRs left out, and repeat this for all subjects. This way the average number of TCRs per fold in the Dash data set varied between 6 (for pp65_123–131_) and 22 (for M45_985-993_), and in the VDJdb dataset between 3 (for p24_30-40_) and 45 (for NS4B_214-222_). The number of subjects and samples for each epitope can be found from Tables 2 and 3.

We also randomly selected a set of background TCRs, so that there was always an equal number of epitope-specific and background TCRs in both training and test sets. We thought this would be the most realistic procedure for the evaluation, as this is likely how these kinds of models will be applied to new data: A model is trained with some set of TCRs and then predictions should be made for TCRs sequenced from an individual from who we have not seen any TCR sequences beforehand.

The data set we gathered from VDJdb contains TCR sequences from multiple studies, many of which have used same conventions for naming their subjects. Therefore we used the combination of the PMID of the publication and the subject id as the subject identifier. For two epitopes, p24_223-231_ and NS3_1406-1415_ there were very few separate subjects, only one and four, respectively. With these epitopes we used 5-fold cross-validation instead of the leave-one-subject-out cross-validation.

### Sequence representation

Computational methods require the data to have some presentation, that they can utilize. Character sequences with variable lengths often provide some challenges as many methods rely on numerical inputs of fixed sizes. One solution is to compare subsequences of same length instead of the complete sequences, which is what for example Generic String kernel (GSkernel) does^42^.

However, by aligning the sequences more approaches become applicable. According to the IMGT definitions^14^ CDR3s can be aligned by introducing a gap in the middle of the sequence (i.e. top of the loop). Alignments for CDR1s, CDR2s, and CDR2.5s can be found from www.imgt.org. When the sequences are aligned, all the sequences within a CDR class (1, 2, 2.5 or 3) have the same length (see Table 1).

We observe sequences *a*_1_*a*_2_ ⋯ *a_L_* of amino acids 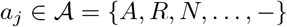 at aligned positions *j* = 1,…, *L*. The alignment guarantees that all sequences have the same possibly padded length *L*. We encode the amino acids *a* with global feature vectors *ϕ*(*a*) ∈ ℝ^*D*^ that associate a *D*-length real-valued code with each of the 21 amino acids including the gap symbol. The sequences are then encoded as data vectors **x** by concatenating the *L* feature vectors into a *D* · *L* length column vectors 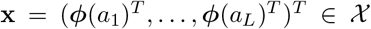. We collect a dataset of *N* sequences into a matrix **X** = (**x**_1_,…, **x**_*N*_)^*T*^ ∈ ℝ^*N*×*DL*^ with rows as sequences and columns as individual amino acid features in aligned order. Each sequence is associated with a class label *y_i_* ∈ {0,1} that indicates whether the sequence was epitope-specific or not. We collect the class labels into an output vector **y** = (*y*_1_,…, *y_N_*)^*T*^ ∈ {0,1}^*N*^.

We can observe amino acid sequences for both the *α*- and *β*-chains and the four complementarity determining regions (CDR) {1, 2, 2.5, 3} from a single TCR. Sequence data for each chain and CDR combination has individual alignments and sequence lengths. We denote the data as (**X**_*α*,1_, **X**_*α*,2_, **X**_*α*,2.5_, **X**_*α*,3_, **X**_*β*,1_, **X**_*β*,2_, **X**_*β*,2.5_, **X**_*β*,3_, **y**).

Substitution matrices such as BLOSUM62^43^ describe the similarity of each amino acid. We modified the BLOSUM62 to include also the gap used in alignments and scaled the matrix values between 0 and 1. The resulting matrix **B** ∈ ℝ^21×21^ is then positive semidefinite. We apply eigendecomposition **B** = **VSV**^*T*^, where the column vectors of **V** encode orthogonal projections of the amino acids on the rows. We use the row vectors of **V**, indexed by the amino acids *a* from the modified BLOSUM62, as our descriptions *ϕ*(*a*) = **V**_*a*_,: with a feature representation *ϕ*(*a*)^*T*^**S***ϕ*(*b*) = [**B**]_*ab*_ for any two amino acids 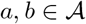. It is possible to use also different substitution models and feature vectors obtained from different sources, or even use the so-called one-hot-encoding, but here we relied only on the eigenvectors of the (gap-extended) BLOSUM62. For our model, we utilized all the 21 components, but in Fig. S8 we show how the amino acids locate on the first two components.

### Gaussian process classification

We use Gaussian process (GP) classification^44^ to predict if a TCR recognizes a certain epitope or not. Gaussian processes model Gaussian distributions of non-parametric and non-linear functions. We apply a link function to squash the function values to a range [0,1] suitable for classification. GPs have a clear advantage of characterizing the prediction uncertainty with class probabilities instead of point predictions. GPs naturally model sequences through kernel functions focusing on sequence similarity as the explaining factor for class predictions.

We use a GP function *f* to predict the latent epitope-specificity *score f*(x) ∈ ℝ of a sequence x. A zero-mean GP prior

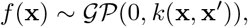

defines a distribution over functions *f*(x) whose mean and covariance are

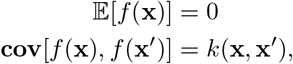

where *k*(·, ·) is the kernel function. We use the standard squared exponential kernel on the vectorized feature representation,

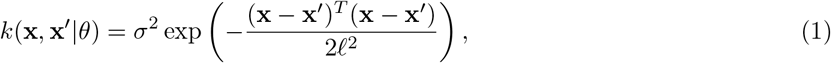

where *ℓ* is the length-scale parameter, *σ*^2^ is the magnitude parameter and *θ* = (*ℓ*, *σ*^2^). For any collection of TCR sequences **X** = (x_1_,…, x_*N*_), the function values follow a multivariate normal distribution

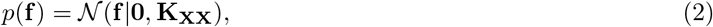

where **f** = (*f*(x_1_),…, *f*(x_*N*_))_*T*_ ∈ ℝ^*N*^ collects all function predictions of the sequences, and **K_XX_** ∈ ℝ^*N*×*N*^ is the sequence similarity matrix with [**K_XX_**]_*ij*_ = *k*(x_*i*_, x_*j*_). The key property of Gaussian processes is that they couple all predictions to be dependent. The Gaussian process predicts similar epitope values *f*(x), *f*(x′) for sequences x, x′ if they are similar according to the kernel *k*(x, x′).

The latent function *f*(x) represents an unbounded real-valued classification score, which we turn into a classification likelihood by the probit link function Φ: ℝ ↦ [0,1],

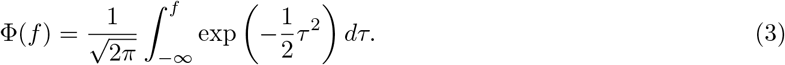

The joint model then decomposes into a factorized Bernoulli likelihood and Gaussian prior,

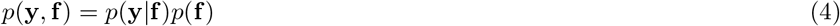

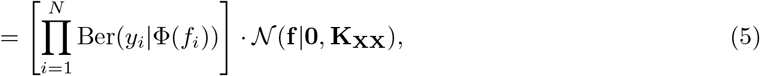

where *f_i_* is a shorthand for *f*(x_*i*_). The objective of Gaussian process modelling is to infer the posterior distribution *p*(**f**|**y**), which is intractable for many non-Gaussian likelihoods. Additionally inferring the kernel hyper-parameters *θ* entails computing the marginalized *evidence*

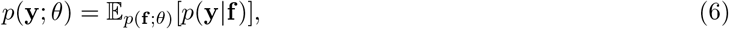

which is also intractable in general and has a limiting cubic complexity 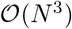^44^. We tackle the scalability with sparse Gaussian processes^45^ and the intractability with stochastic variational inference^46^.

### Variational inference for low-rank GP approximation

We consider low-rank sparse Gaussian processes by augmenting the system with *M* inducing *landmark* pseudosequences 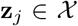 with associated (label) function values *u_j_* = *f*(**z**_*j*_) ∈ ℝ. We collect all inducing points into structures **Z** = (**z**_1_,…, **z**_*M*_)^*T*^ and **u** = (*u*_1_,…, *u_M_*)^*T*^. By conditioning the GP with these values we obtain the augmented Gaussian process joint model

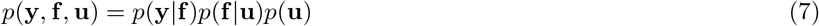

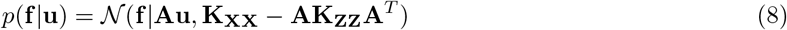

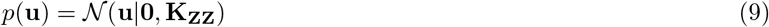

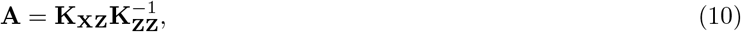

where **K_XX_** ∈ ℝ^*N*×*N*^ is the kernel between observed sequences, **K_XZ_** is between observed and induced sequences and **K_ZZ_** is between induced sequences. The matrix **A** projects the *M* inducing points to the full observation space of *N* sequences.

Next, we define a variational approximation for the inducing points,

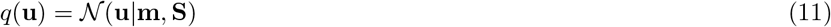

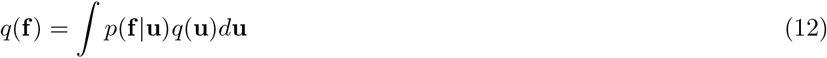

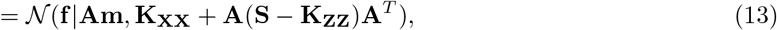

where **m** ∈ ℝ^*M*^ and **S** ≽ **0** ∈ ℝ^*M*×*M*^ are free variational parameters to be optimized. It can be shown that minimizing the Kullback-Leibler divergence KL[*q*(**u**)||*p*(**u**|**y**)] between the approximative posterior *q*(**u**) and the true low-rank posterior *p*(**u**|**y**) is equivalent to maximizing the evidence lower bound (ELBO)^47^

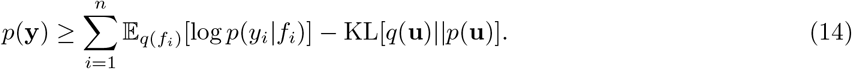

The log expectation is tractable for Probit likelihoods^48^, while the KL term similarly has a closed form for two Gaussian densities.

Due to the small data regime we choose the optimal assignment of selecting **Z** = **X** and **u** = **y**, which corresponds to the full Gaussian variational approximation of Nickish et *al*.^49^, while for larger datasets the inducing landmark points can also be optimised^46^. We then optimize the ELBO (14) with respect to the variational parameters **m** and **S** as well as the kernel hyperparameters *θ*, that is, the lengthscales *ℓ_cr_* and weights *w_cr_*.

Finally, predictions **f**_*_ of new test sequences 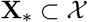 follow a variational predictive posterior

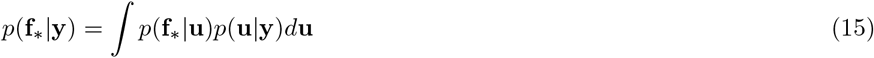

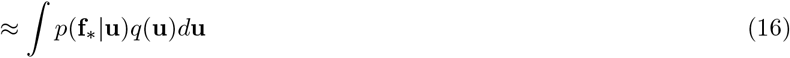

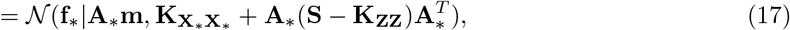

where **A**_*_ indicates projection from the landmark points **Z** to the new sequences **X**_*_. The predictive distribution is a Gaussian distribution for the latent test values **f**_*_, from which the distributions of the test labels can be retrieved through the link function.

We have implemented our model using GPflows VGP-model^50,51^. Our code, data sets, and some examples can be found from github.com/emmijokinen/TCRGP.

### Multiple kernel learning

As mentioned in Section 4.1, when a TCR binds to a pMHC, its CDR3*β* is presumably always in contact with the peptide while the other CDRs may contact the peptide, but mainly contact the peptide binding groove of the MHC presenting the peptide.^13^ took this into account by giving fixed weights for the distances between amino acids within different CDRs, giving more weight to the CDR3. As it can vary which CDRs can be in contact with different peptides, we did not want to determine the importance of these different CDRs beforehand, but instead created separate kernels for each CDR and let our model decide which of them are important. We define the kernel as a convex combination of the four CDR regions *r* and the two chains *c*,

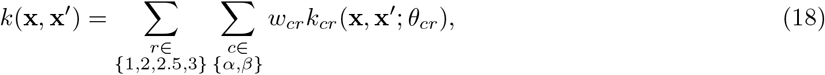

where the weights *w_cr_* ≥ 0 are non-negative.

### TCR repertoire diversity

To estimate the diversity of the epitope-specific TCRs for each epitope, we developed a diversity measure following the example of Dash et *al*.^13^. The Simpson’s diversity index was then generalized to account for the similarity of TCRs by utilizing the Gaussian kernel function as follows:

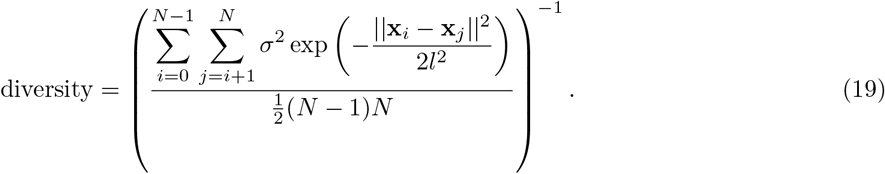

Here *σ*^2^ is the kernel variance and *l* is the lengthscale of the Gaussian kernel used by TCRGP, and x_*i*_ and x_*j*_ are feature vectors for the TCRs *i,j* ∈ [1, *N*]. The kernel variance and lengthscale were set to the average values used for the 22 epitopes in the VDJdata (*σ*^2^ = 5.52, *l* = 2.50).

### TCRGP classifiers for HBV-epitopes

Utilizing the TCRs specific to HBV epitopes HBV_core169_, HBV_core195_, HBV_pol282_, and HBV_pol387_ from Cheng et *al*.^29^ and control sequences from Dash et *al*.^13^ we trained a TCRGP classifier for each epitope. We utilized all epitope-specific TCRs from which we could determine also CDR1*β*, CDR2*β*, and CDR2.5*β* in addition to CDR3*β* and complemented these epitope-specific TCRs with the same amount of control TCRs. We considered TCRs which were predicted to recognize the epitopes with 85 % probability as epitope-specific. The amounts of epitope-specific TCRs and AUROC-scores obtained from leave-one-subject-out cross-validations for each epitope are shown in Table 4. We used TCRGP and VGP with all epitopes except for HBV_pol387_, with which we used SVGP with 700 inducing points due to the high number of samples.

**Table 4:** HBV-epitopes for which we trained TCRGP classifiers, the numbers of epitope-specific TCRs and subjects and mean AUROC-scores from leave-one-subject-out cross-validations.

| Epitope | Samples | Subjects | AUROC |
| --- | --- | --- | --- |
| HBV<sub>core169</sub> | 699 | 9 | 0.756 |
| HBV<sub>core195</sub> | 588 | 12 | 0.847 |
| HBV<sub>pol282</sub> | 459 | 12 | 0.880 |
| HBV<sub>pol387</sub> | 1348 | 12 | 0.760 |

### Single-cell RNA-sequencing analysis

The unnormalized expression count data of T cells passing the quality control in the Zheng data were fetched from GEO (GSE98638) along with the TCR*αβ*-sequences inferred from the full-transcript single-cell RNA-sequencing data and inferred phenotypic states as described by Zheng et *al*.^28^. As the TCR*β*-sequenced training data for HBV-specific epitopes was HLA-A-restricted, we focused our analysis only on T cells capable of peptide recognition in HLA-A restricted manner, namely clusters CD8-LEF1, CD8-CX3CR1, CD8-LAYN and CD8-GZMK. The data was log-normalized to 10 000 counts per cell and scaled accordingly with the Seurat 3.0.2.^52^ package for R 3.5.2. The highly variable genes (HVGs) were chosen as the genes showing the highest mean to variance ratio (min expression = 0.5, max expression 3, min variance 0.5) with the FindVariableFeatures-function. The linear dimensionality reduction was calculated with PCA for the scaled expression matrix containing only HVGs. Non-linear dimensionality reduction was performed with UMAP for principal components that had standard deviation > 2 using standard parameters with the RunUMAP-function. To receive a better grouping for the selected cells, we used a graph-based clustering approach implemented in the Seurat tool. To find the shared nearest neighbor graph, the function FindNeighbors was used with the same amount of PCs as with UMAP. To determine optimal clustering, FindClusters-functions was used with several parameter values for the resolution parameter, ranging from [0.1, 3]. The optimal clustering was decided by agreement of grouping in the UMAP-embedding and the labels from clustering by visual interpretation. The cytotoxic and exhaustion signatures for the clusters were calculated as cell-wise mean expression of cytotoxic (*NKG7, CCL4, CST7, PRF1, GZMA, GZMB, IFNG, CCL3*) and exhaustion genes (*CTLA4, PDCD1, HAVCR2, TIGIT, LAG3*). The one-sided Fisher’s test for enrichment of epitope-specific T cells to different phenotypes was calculated independently for individual and pooled patients, epitopes and tissues which were then adjusted with Benjamini-Hochberg for false-discovery.

## Data and code availability

TCRGP software tool and the data sets used for the evaluation of the method are available at https://github.com/emmijokinen/TCRGP. Software and data for the single-cell RNA-sequencing analysis of HCC-patients are available at https://github.com/janihuuh/tcrgp_manu_hcc.

## Acknowledgements

We would like to acknowledge the computational resources provided by the Aalto Science-IT. This work has been supported by The European Research Council (M-IMM project), Academy of Finland (project numbers: 287224, 299915, 313271, 314442 and 314445), Finnish special governmental subsidy for health sciences, research and training, the Sigrid Juselius Foundation and the Finnish Cancer Societies

## Author Contributions

All authors contributed to designing the study. E.J., M.H. and H.L. co-developed the method. E.J. implemented TCRGP and carried out the experiments with support from other authors. J.H. developed the analysis approach for scRNA+TCR*αβ*-sequencing data from HCC-patients and performed the analysis with support from other authors. S.M. helped with analyzing the results. All authors contributed to the manuscript writing.

