## Supplementary material for "Determining epitope specificity of T cell receptors with TCRGP"

### Determining epitope specificity of T cell receptors with TCRGP: Supplementary material

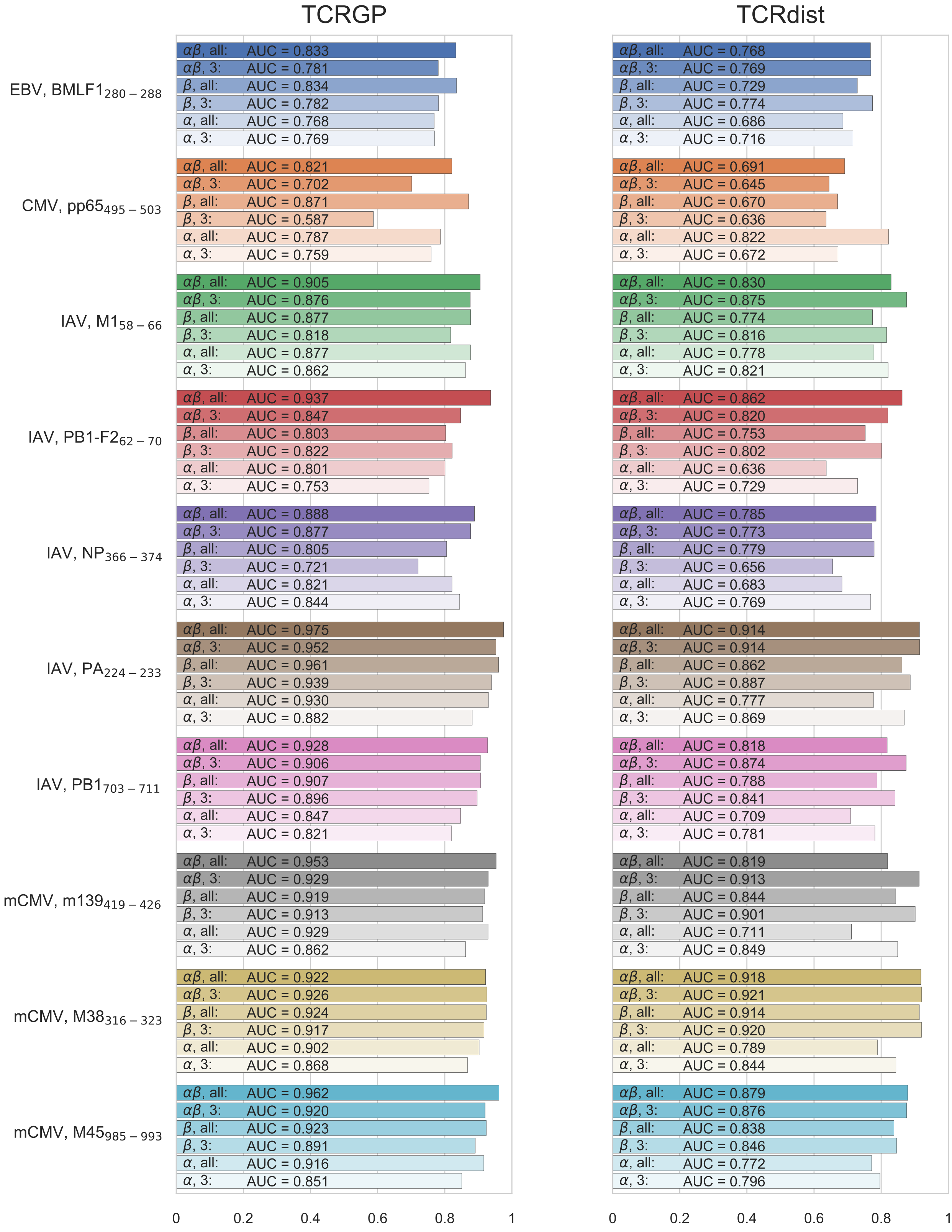


Figure S1: **Mean AUROC scores for the Dash data using leave-one-subject-out cross-validation**. TCRGP models (left column) and TCRdist models (right column) we trained using either only CDR3 or all CDRs from TCR$\alpha$, TCR$\beta$, or both and either with.

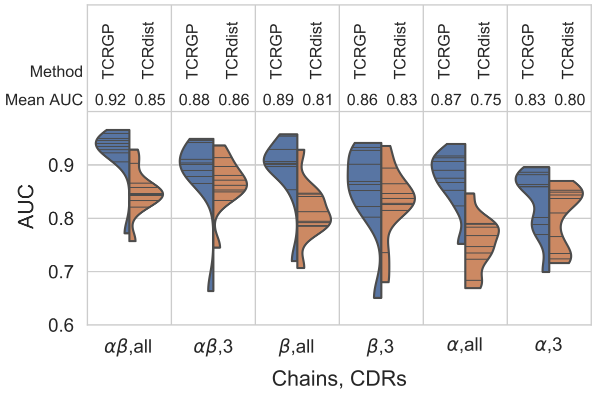


Figure S2: **Mean AUROC scores for the Dash data using leave-one-out cross-validation.** Only unique TCRs have been utilized. The blue parts of the violin plots illustrate the AUC-scores of predictions made by TCRGP for all the epitopes. The orange sides illustrate the AUC-scores obtained with TCRdist. A horizontal line within a violin plot presents the mean AUC-score obtained for one epitope. The used chains ($\alpha$ and/or $\beta$) and CDRs (three or all) are indicated below each panel.

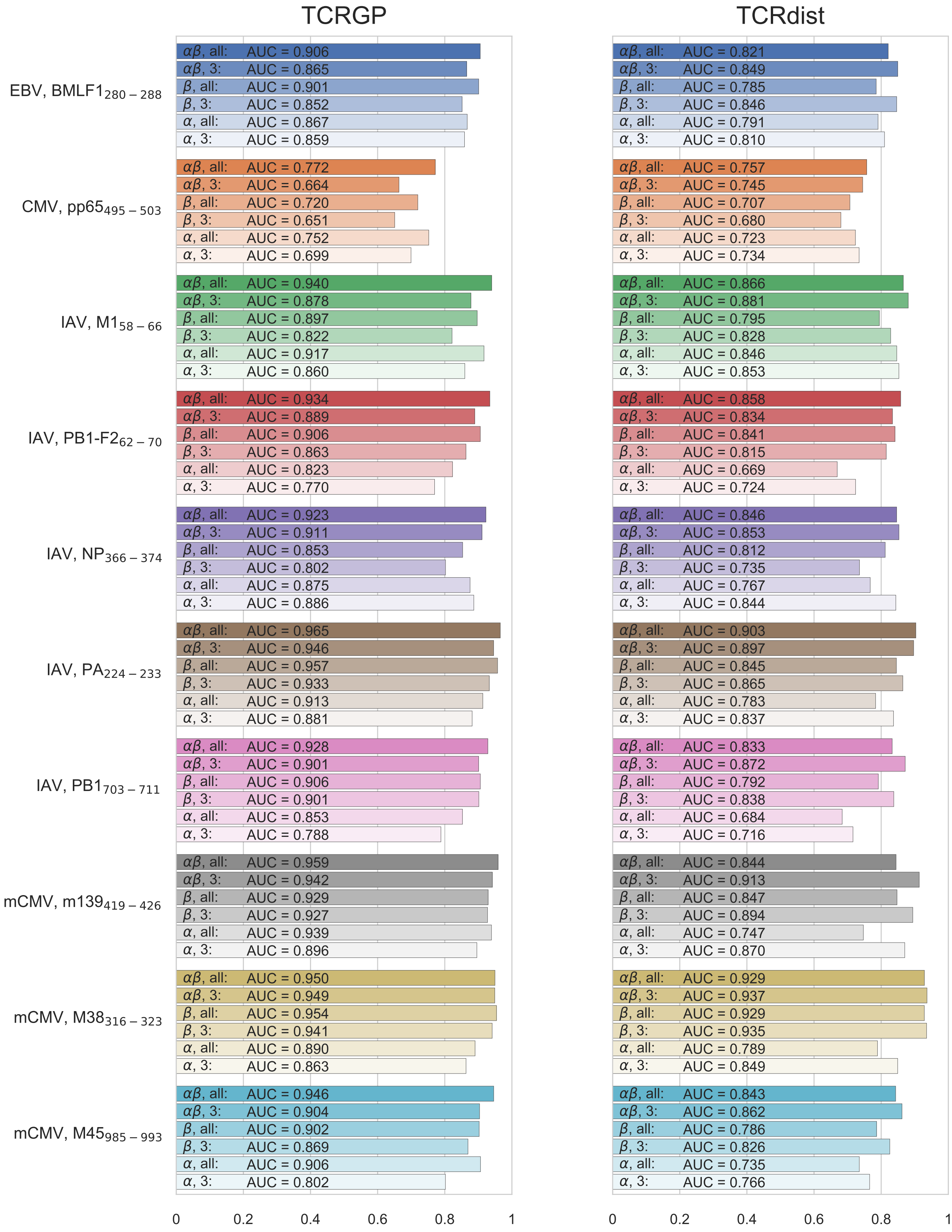


Figure S3: **Mean AUC scores for the Dash data using leave-one-out cross-validation.** Only unique TCRs have been uilized. TCRGP models (left column) and TCRdist models (right column) we trained using either TCR$\alpha$, TCR$\beta$, or both and either with only CDR3 or all CDRs.

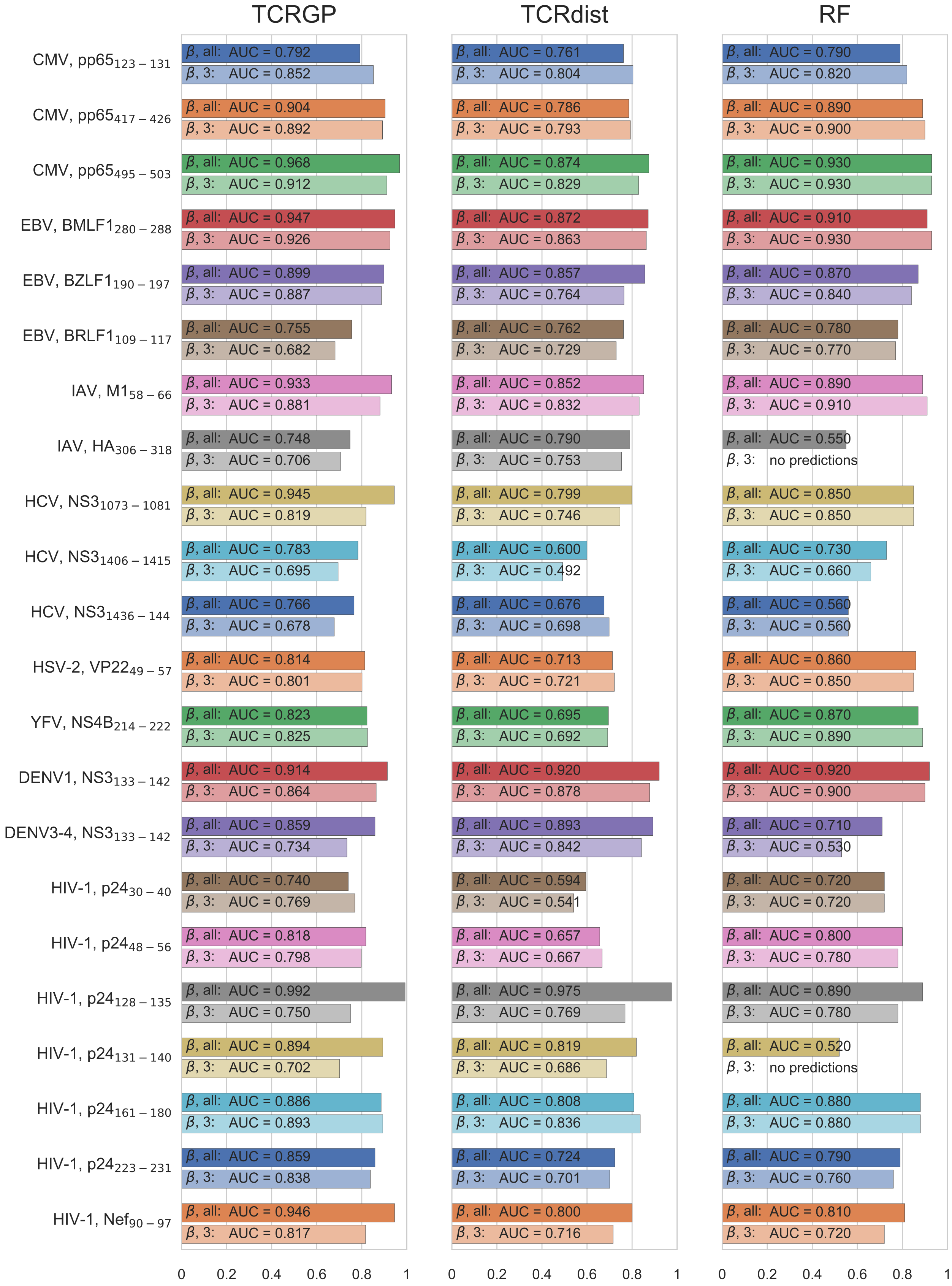


Figure S4: **Mean AUROC scores for the VDJdb data using leave-one-subject-out cross-validation.** TCRGP models (left column) and TCRdist models (middle column) were trained using TCR$\beta$ with either only CDR3 or all CDRs. TCR-classifiers by De Neuter *et al.*[1] (right column) were trained using the CDR3$\beta$ and the V$\beta$-gene, from which the other CDRs can be derived from.

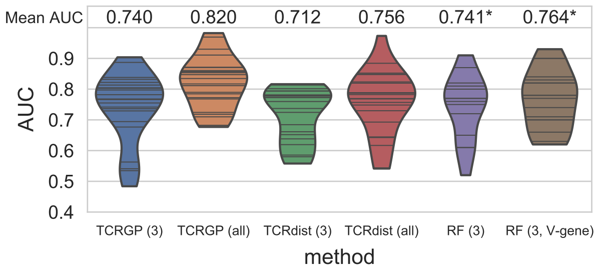


Figure S5: **Mean AUROC scores for the VDJdb data using leave-one-out cross-validation.** Only unique TCRs have been utilized. One violin plot presents the mean AUROC-scores obtained with one method for all epitopes in our VDJdb data. Below each violin plot there is the name of the method used and in the brackets which CDRs have been used (3 for CDR3, all for CDR1, CDR2, CDR2.5, and CDR3). A horizontal line within a violin plot presents the mean AUROC-score obtained for one epitope. RF (Random Forest TCR-classifier of De Neuter *et al.*[1]) has been left out as it could not produce predictions for all epitopes. Results for individual epitopes of the VDJdb data are shown in Fig. S6.

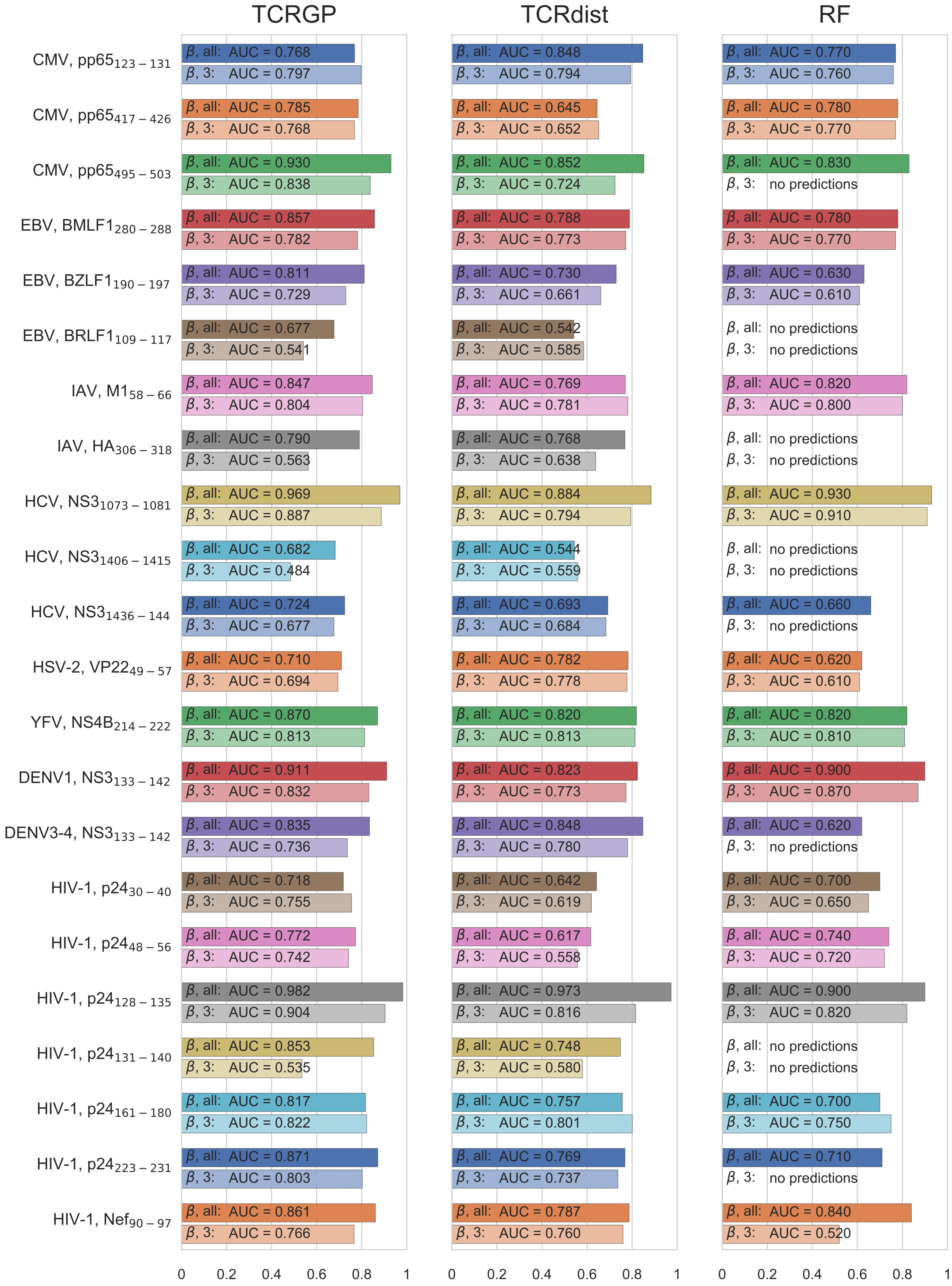


Figure S6: **Mean AUROC scores for the VDJdb data using leave-one-out cross-validation.** Only unique TCRs have been utilized. TCRGP models (left column) and TCRdist models (middle column) were trained using TCR$\beta$ with either only CDR3 or all CDRs. TCR-classifiers by De Neuter *et al.*[1] (right column) were trained using the CDR3$\beta$ and the V$\beta$-gene, from which the other CDRs can be derived from.

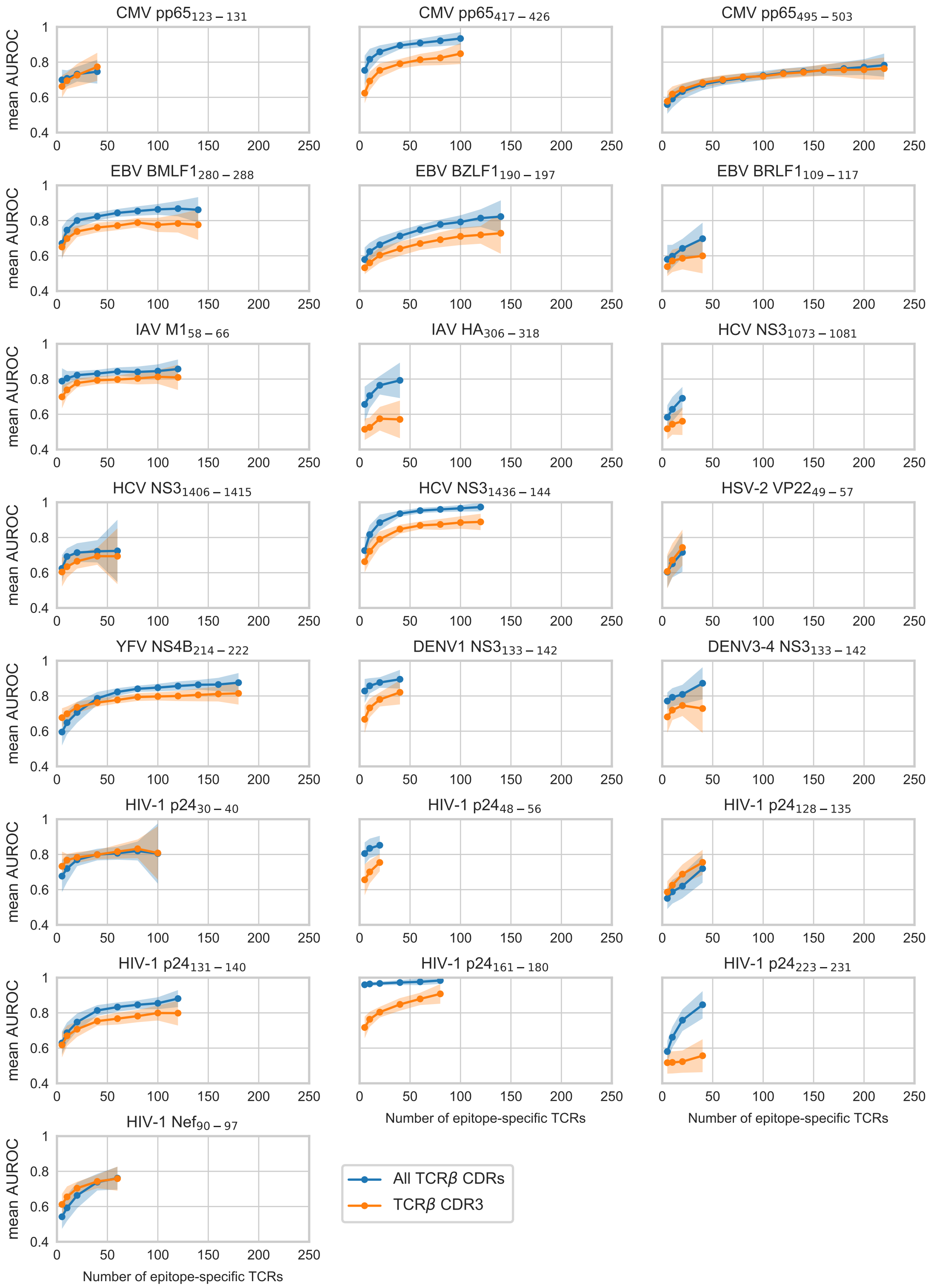


Figure S7: **Learning curves**. With each epitope from the VDJdb dataset, TCRGP models were trained using different numbers of unique epitope-specific TCRs, always complemented with the same number of control TCRs. For each point of the learning curve the model was trained with 100 random samples of the TCRs, using either CDR1, CDR2, CDR2.5, and CDR3 (blue curves), or only CDR3 (orange curves). The darker lines show the mean of the predictions and the shaded areas +/- the standard deviation for the 100 folds.

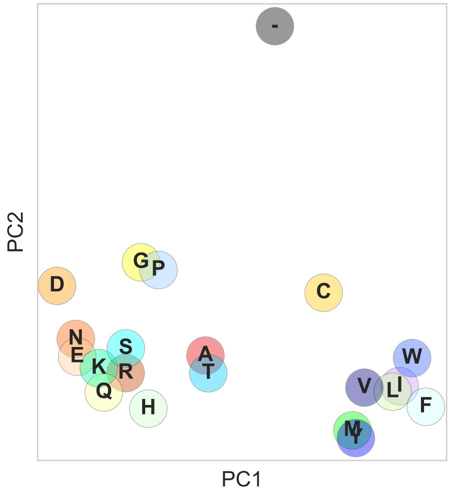


Figure S8: **PCA components of BLOSUM62.** All 20 amino acids and the gap located on the first two orthogonal components of the modified BLOSUM62 substitution matrix. The amino acids are coloured according to Ruiz and Lefranc [2].
